# Effects of kin recognition on root traits of wheat germplasm over 100 years of breeding

**DOI:** 10.1101/2020.09.04.243758

**Authors:** Lars Pødenphant Kiær, Jacob Weiner, Camilla Ruø Rasmussen

## Abstract

Plant root and shoot growth has been shown to depend on the relatedness of co-cultivated genotypes, implying the existence of ‘kin recognition’ mechanisms mediated by root exudates. If confirmed, this has important implications for crop breeding.

We present the first large-sale investigation of kin recognition in a crop germplasm collection comprising 30 North-European cultivars and landraces of spring wheat, spanning 100 years of breeding history. In a full diallel *in vitro* bioassay, we compared root growth of seedlings when growing in pure substrate, or in substrate previously occupied by a donor seedling from the same (KIN) or another (NONKIN) genotype.

Seedlings growing in KIN or NONKIN substrate generally had longer but not more roots than seedlings growing in pure substrate. Responses were generally larger in longer roots, suggesting that root elongation was promoted throughout the growth period. Responses to KIN and NONKIN substrates were found to range from positive to negative, with root length responses to kin being increasingly positive with year of release. Seedlings growing in KIN substrate generally had shorter but not fewer roots than seedlings growing in NONKIN substrate. This kin recognition ranged from positive to negative across the specific donor-receiver combinations and did not change systematically with year of release of either genotype. Root traits in both KIN and NONKIN substrate were affected by both donor and receiver genotype, and these effects were generally larger than the effect of specific combinations. Genotypes showing higher levels of kin recognition also tended to invoke larger responses in other genotypes. Kin recognition was reduced in most cases by the addition of sodiumorthovanadate, a chemical inhibitor, supporting the hypothesis that kin responses were mediated by changes in the chemical constitution of the substrate.

The identified patterns of kin recognition across the germplasm collection were complex, suggesting a multigenic background and shared breeding history of the genotypes. We conclude that kin response represents a potential target for crop breeding which can improve root foraging and competitive interactions.

## Introduction

As sessile organisms, plants have evolved a wide range of mechanisms that allow individuals to adapt continuously to their environment and maximize their growth, survival, and reproductive success. Plasticity of plant traits in response to the many chemical, physical and biological cues in the soil environment have thus been found to promote complex, integrated developmental trajectories, including nutrient foraging, competition with other plant species, and investment in promoting specific beneficial microorganisms.

A growing number of studies have demonstrated the ability of plants to distinguish their own roots from those of neighbouring plants. There is also evidence that some plants are able to distinguish closely related neighbours (kin) from more distant relatives, resulting in plastic changes that limit “selfish” root proliferation and alter allometric relationships such as allocation to roots and shoots (Dudley and File 2007, Murphy and Dudley 2009, Biedrzycki et al. 2010, Biernaskie 2011, Bhatt et al. 2011, Crepy & Casal, 2015), overall plant growth (Marler 2013) and morphology (Biedrzycki et al. 2010; Semchenko et al. 2014; Crepy and Casal 2015), allocation to reproduction (Donohue 2003, Biernaskie 2011), and spatial orientation of roots (Fang et al. 2013).

The patterns of kin recognition behaviour in plants are not well described, and the direction and extent of kin recognition seems to differ among plant groups, ranging from more aggressive to more evasive root growth in the presence of nonkin. Some studies have failed to find evidence for kin recognition (Argyres & Schmitt 1992, Dudley & File 2007, Monzeglio & Stoll 2008, Milla et al. 2009, Murphy & Dudley 2009, Masclaux et al. 2010), suggesting that it is not consistently expressed or that it may be less important than other ecological interactions such as competition (Masclaux et al. 2010). Studies have found kin response to be moderated by environmental factors such as plant density (Lepik et al. 2012), nutrient availability (Sattler and Bartelheimer 2018, Li et al. 2018) and heavy metal concentration in the soil (Li et al. 2018).

These previous findings indicate that the genetic background and evolutionary role of kin recognition in plants may be complex. The mechanisms behind it are not elucidated but results to date suggest that information on neighbour identity comes from root exudates (Biedrzycki et al. 2010) and involve biochemical pathways related to plant defence in *Arabidopsis thaliana* (Biedrzycki et al. 2011a).

Behaviour informed by kin recognition is hypothesized to help individuals avoid costly competition with close relatives. Helping a close relative increases the fitness of the altruist indirectly, a concept called kin selection (Hamilton 1964). It has also been hypothesized that plant phenotypic responses to neighbours, such as shade avoidance and root proliferation in response to neighbours, are advantageous for individuals but detrimental at population level (Weiner 2004). If plants can distinguish between closely and distantly related neighbours and behave differently, it could have important implications for plant evolution. And if this ability exists in crop plants, it could play an important role in increasing yields and/or resource use efficiency in plant production (Bais 2015).

Some crop species have been found to proliferate roots in response to neighbouring roots (e.g. Zhu et al. 2019), but in many cases this may be a response to reduced nutrient levels, not neighbouring roots *per se* (McMickle and Brown 2014). A study used unfertilized transparent gel to show that roots of rice tended to avoid neighbouring root systems of plants of a different genotype, but not of the same genotype (Fang et al. 2013). While suggesting the existence of nutrient-independent root-root mediated kin response in a cereal crop, the direction of the response seems contrary to the hypothesized competition avoidance among kin. Inbreeding cereal crops such as wheat are predominantly grown as monocultures, in which all individuals are bred and propagated to be as uniform and closely related as possible, conforming to the definition of *kin*. It remains unknown if breeding has affected kin recognition ability during cereal domestication, particularly in light of the intensive breeding during the 20^th^ century leading to increasingly homogeneous cultivars.

We present here the first large-sale investigation of kin recognition in a crop germplasm collection, and the first in bread wheat (*Triticum aestivum*). We use a screening bioassay to test the hypotheses that (1) kin recognition behaviour is found in wheat already in the earliest growth stages, (2) wheat roots generally grow shorter when exposed to kin as compared to nonkin growth substrate, in accordance with kin selection theory, (3) this is due to changes in the chemical composition of the substrate, and (4) kin recognition behaviour has been reduced by the intensified monoculture breeding throughout the 20^th^ century.

## Materials and Methods

### Genetic material

Seeds from 30 North-European genotypes of bread wheat (*Triticum aestivum*) were obtained from seedbank repositories (NordGen, Gatersleben IPK). These represented germplasm from 100 years of breeding (Table S1), with 24 genotypes being cultivars released in the period 1900 to 1997 and six landraces being of undefined pre-1900 origin. The 20 most recent cultivars were selected among a larger set of 50 cultivars evaluated for genetic variation based on SSR markers in the context of another study (LP Kiær, unpublished), being among the cultivars with the highest level of genetic purity. All genotypes were propagated in greenhouse pots and field plots, following vernalization of winter types (see Table S1), and their seeds were harvested, threshed, and stored for further testing.

### Bioassay

Seedlings of each genotype were grown in a water agar substrate made of 3g Agargel^™^ (Sigma-Aldrich Co. LLC) per 1000ml deionized water with no nutrients added, mixed in a magnetic stirrer and sterilized in an autoclave (reaching 121°C for 15 min). Upon cooling to approx. 40°C, 3ml water agar was transferred to each well of a VWR 12-well cell culture multiplate (flat bottom, non-treated), using a BRAND seripettor^®^ pro dispenser in a laminar flow cabinet to reduce the risk of contamination. Multiwell plates were then incubated in a Binder KBW 400, using a cycle of 14h day (4500 lux) at 22°C and 10h night (dark) at 14°C.

Unsterilized seeds were pre-germinated in the dark on moist filter paper in Petri dishes. After approximately 48 hours, individual seedlings were positioned carefully in a well with rootlets (hereafter ‘roots’), covered with substrate, using sterilized tweezers in a laminar flow cabinet. Fungal infection was observed in only very few samples, which were discarded. Only seeds with normal germination and growth were assessed and analysed.

A full diallel bioassay design was used, exposing seedlings of each genotype, as *receivers*, to a growth substrate that was previously occupied by another *donor* seedling from the same (KIN) or another (NONKIN) genotype, for a total of 900 genotype combinations. In one replicate of a given combination (placed in one multiplate well), a seedling of the *donor* genotype was grown in the incubator for a period of six days and then removed, carefully leaving all substrate in the well. A newly germinated seedling of the *receiver* genotype was then placed in the same well and grown in the incubator for another period of six days, and then removed for further root trait assessment (see below). A subset of seedlings from the first growth period were sampled for further root trait assessment, providing a reference treatment in pristine substrate without exposure to other seedlings than the individual itself (PURE). The average number of replicates were 12.8 for KIN treatments, 2.5 for NONKIN treatments and 10.1 for PURE treatments.

To test the hypothesis that KIN and NONKIN responses were attributable to organic chemicals released to the substrate by the previous genotype, the 60 most responsive genotype combinations, and the corresponding KIN treatments of *receiver* genotypes, were grown with (*inhib*) or without (*control*) added sodium orthovanadate (Na_3_VO_4_). This is an alkaline phosphatase known to act as an inhibitor of several enzyme classes and other organic compounds. The inhibitor was added to the water agar substrate in the cooling phase following sterilization, to a final concentration of 150μM. The number of replicates was between 4 and 5, with an average of 4.8 for KIN_*control*_ and KIN_*inhib*_ treatments, 4.4 for NONKIN_*control*_ treatments and 4.5 for NONKIN*inhib* treatments.

### Root trait assessment

Roots from each removed seedling were cut manually at the seed base and mounted individually under a plastic sheet before scanning on a flatbed scanner at 600 dpi resolution. Scanned images (Fig. S1in Supplementary) were analysed in Matlab, using proprietary code (available on request), giving data on number of roots, length of individual roots and average root width. Samples with three roots or fewer were discarded to avoid influence of any seedlings not growing well (2.7% of the full diallel samples, and 0.5% of the inhibitor bioassay samples). Most seedlings produced at least five roots, in which case the five longest roots were considered as the higher-ranking primary root (P) and the first (F) and second (S) pairs of seminal roots.

Six root traits were used to assess root growth and kin recognition. The number of roots (RN) was used as a measure of root initiation, independent of individual root lengths. Length of the longest root (RL-MAX) was used as a measure of root growth potential. The total root length (RL-TOTAL) was derived as the summed length of all roots, and is considered a measure of total root activity. The coefficient of variation of seedling root lengths (RL-CV) was used as an overall measure of root uniformity. The summed length of P, F and S roots (RL-PFS) was used as a measure of primary root growth, in cases where at least five roots were observed. Total root volume (RV) was used as a proxy for root biomass, considering roots as tubes of a given average width (RW), i.e. RV = π · (RW/2)^2^ · RL-TOTAL.

### Calculation of kin and nonkin responses and effects

Root traits were analysed within the response-and-effect framework developed in the context of trait-based ecology (Garnier et al. 2015), considering any effects and responses as indirect interactions via the substrate environment (Fig. 1).

**Figure 1.**
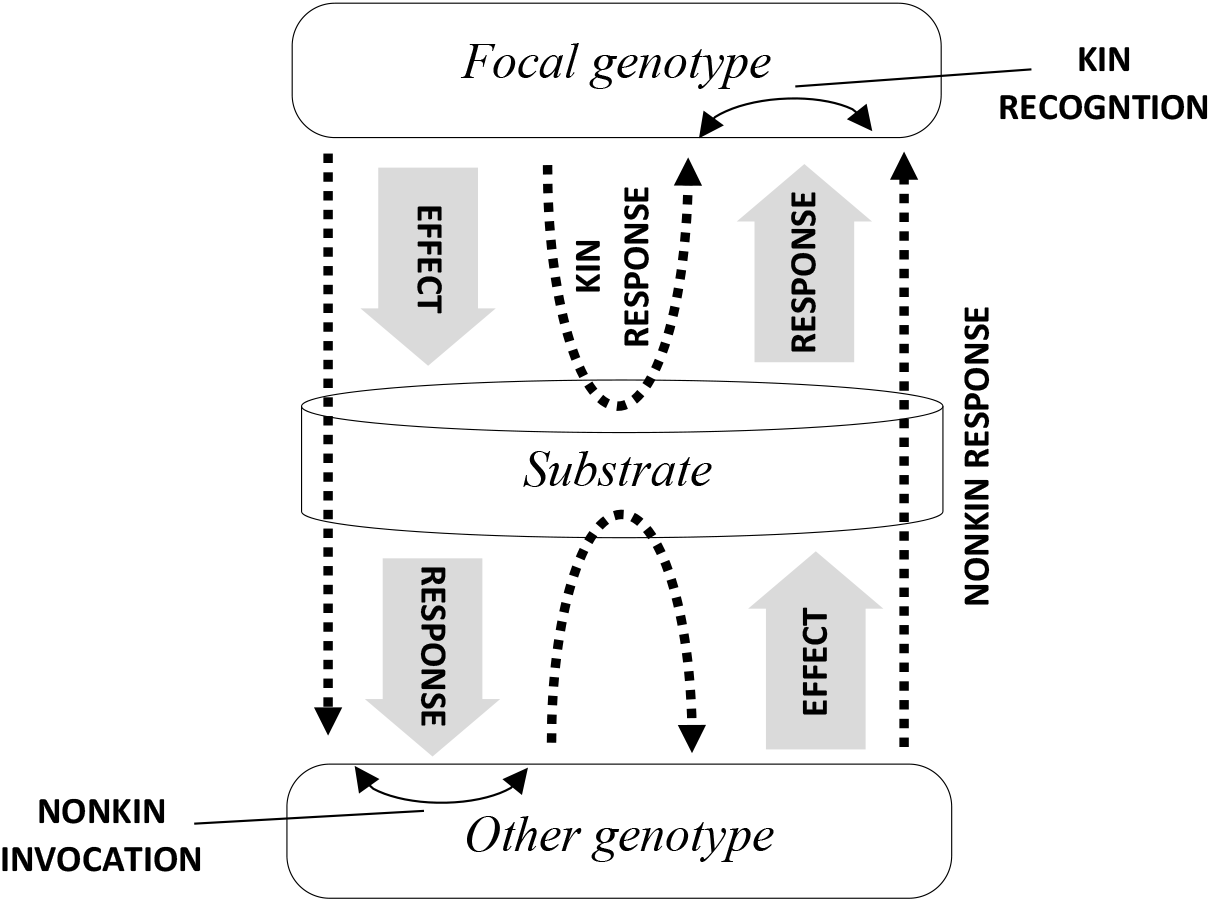
Model showing the indirect responses and effects between a focal genotype and another genotype via a shared substrate environment. Also shown are routes of kin response and nonkin response of the focal genotype and similar responses of the other genotype (dotted lines), as compared to growth in pristine substrate (PURE). The identification of kin differentiation and nonkin invocation through comparison of these responses is shown as double arrows.

Basic root growth of each genotype was identified based on the root traits of seedlings growing in pristine substrate (PURE). *Kin response* was defined as the change in a root trait of a focal genotype when growing in substrate following a *donor* seedling from the same genotype (KIN) compared to basic root growth in the PURE treatment. Overall kin response was calculated for each focal genotype as the average change across donors and replicates. *Nonkin response* was similarly defined as the change in a root trait of a focal genotype when growing in substrate following a seedling from another genotype (NONKIN) compared to basic root growth in the PURE treatment. Overall nonkin response was calculated for each focal genotype as the average change across all replicated NONKIN treatments of that genotype.

*Kin recognition* was defined for a given *donor-receiver* genotype pair as the change in a root trait of the *receiver* genotype when growing in KIN substrate compared to growing in NONKIN substrate (following the *donor*). Overall kin recognition was calculated for each focal genotype as the average kin recognition across all replicated NONKIN treatments of that genotype.

*Nonkin invocation* was defined for a given donor-receiver genotype pair as the root response invoked by the focal genotype (as *donor*) in the other genotype (as *receiver*) as compared to that other genotype growing in its corresponding KIN substrate. Overall nonkin invocation was calculated for each focal genotype as the average of all replicated nonkin responses it invoked in other genotypes.

### Statistical analysis

Data were analyzed with R (version 4.0.1, R Core Team 2020), using core functions unless otherwise specified.

Basic root traits of genotypes were estimated based on the assessment of seedlings grown in PURE substrate. Pairwise correlations among root traits were tested using Pearson’s product moment correlation. For each trait separately, a linear model with *genotype* as independent variable was then used to obtain genotype-specific estimates and test for overall differences between genotypes, using one-way ANOVA. Root volume was square root transformed before analysis to achieve normality. Correlation between root traits and the year of release (excluding landraces) were tested using Pearson’s product-moment correlation. To include the landraces, which have no release year, in additional correlation analyses, they were assigned a release year immediately prior to the earliest cultivar genotype (i.e. 1895-1900).

Overall changes in root traits when exposed to KIN substrate were tested based on the combined KIN and PURE dataset, using t-test of the effect of treatment (KIN or PURE) in a linear model, with a subset of models including *receiver genotype* as covariate or the *relatedness x genotype* interaction. Effects of NONKIN (compared to PURE) substrate on root traits were tested using the same approach. Kin recognition and nonkin invocation were analysed using t-test of the effect of *relatedness* (KIN or NONKIN) in a linear model, with a subset of models including *receiver genotype* as covariate or the *relatedness x genotype* interaction. For these and other tests of effect on RN, a zero-truncated negative binomial model was used, as implemented in the R package *VGAM* (Yee 2020).

To quantify the effect of *donors* (*d*) and *receivers* (*r*) on root traits in the full diallel setup, we applied the concept of combining ability (Sprague & Tatum 1942). Here, general combining ability (GCA) is defined as the average performance of a genotype in a series of combinations with other lines, and specific combining ability (SCA) is the effect of interaction between specific genotype pairs. Griffing’s model III with reciprocals and random effects, as implemented in the R package *DiallelAnalysisR* (Yaseen 2016), was used to estimate general and specific *donor-receiver* effects for each root trait. This was not estimated for RL-PFS because of missing values in some combinations.

Estimates of genotype-specific kin recognition and nonkin invocation were derived for each root trait using one-way ANOVA, and correlations between kin recognition and nonkin invocation were tested for each root trait using Pearson’s product-moment correlation. The effect of chemical inhibitor on genotype-specific kin recognition was tested for each root trait, using one-way ANOVA.

## Results

### Basic root growth of genotypes

Root traits of seedlings tested in the PURE treatment showed significant genotypic variation (Table 1; Table S2). RN was less variable, with most individuals producing from 4 to 7 roots. A few individuals produced up to 10 roots, of which the lower-ranking roots were typically very short (not shown). Genotypes accounted for most of the variation in root traits (*R*^2^-values between 0.92 and 0.99). Length-related root traits, i.e. RL-MAX, RL-PFS and RL-TOTAL were positively correlated, both with and without genotype as a cofactor (not shown).

**Table 1.** Analysis of variance of effects of genotype and year of release, respectively. Landraces are included in both analyses. Slopes from linear regression of each root trait against year of release are provided. ns, *, **, *** denote non-significance and significance at P < 0.05, P < 0.01 and P < 0.001, respectively.

| Root trait | Difference between genotypes? | Correlation with year of release? |
| --- | --- | --- |
| RL-MAX | *** | 0.1047 *** |
| RL-PFS | *** | 0.3174 *** |
| RL-TOTAL | *** | 0.3447 *** |
| RL-CV | ** | 0.0001 * |
| RV | *** | 0.0124 *** |
| RN | ns | ns |

Root length generally increased with the year of release (Table 1). The number of roots did not increase, suggesting that this was mainly due faster root elongation. While considerable variation was seen around regression lines (Fig. S2), the regressions reveal that the cultivar *Saffran* (from 1978) had markedly lower root volume than expected from its year of release, whereas the landrace *Lantvete från Halland* had markedly higher root volume than expected (Fig. S2e).

### Kin and nonkin responses

Seedlings from the KIN treatment generally had higher root growth rates than seedlings from the PURE treatment (Table 2). This effect was strongest for longer roots, resulting also in a higher RL-CV. There was no significant effect on RN (Table 2). Kin response did not differ significantly among genotypes, i.e. the interaction term *relatedness x genotype* was not significant for any root trait (not shown). Genotypic kin responses in RL-TOTAL and RL-PFS increased with year of release from mainly negative to mainly positive (both with P < 0.05).

**Table 2.** Overall kin responses, nonkin responses and kin recognition for each root trait, using two-way ANOVA accounting for *genotype* and *relatedness*. Separate analyses where made for each root trait. Percentages were calculated from the main effects, with the shown RV responses being based on the untransformed values. ns, *, **, *** denote non-significance and significance at P < 0.05, P < 0.01 and P < 0.001, respectively.

| Root trait | Kin response (%) | Nonkin response (%) | Kin recognition (%) |
| --- | --- | --- | --- |
| RL-MAX | +8.0 *** | +11.8 *** | -3.4 * |
| RL-PFS | +4.2 ** | +7.5 *** | -2.8 * |
| RL-TOTAL | +2.8 * | +6.2 *** | -3.2 * |
| RL-CV | +22.7 *** | +18.6 *** | +3.5 (*) |
| RV | +25.2 *** | +29.6 *** | -3.5 (*) |
| RN | +1.9 ns | -0.2 ns | +2.1 ns |

Seedlings from the NONKIN treatment generally had significantly higher root growth rates compared to seedlings from the PURE treatment (Table 2). This was more pronounced for the longer roots, matched by higher RL-CV in NONKIN treatments (Table 2). There was no effect on RN. Nonkin responses did not differ significantly among genotypes, i.e. the interaction term *relatedness x genotype* was not significant for any root trait (not shown). RV response tended to decrease with year of release, as seen from a marginally significant interaction term (*relatedness* x *year*; P = 0.055), suggesting that positive nonkin responses in root volume were generally more common in genotypes with earlier release date.

All kin and nonkin responses and effects varied substantially among genotypes, ranging from positive to negative (Table 3). We did not find correlations between kin or nonkin responses and measurements in the PURE treatment for any of the root traits (not shown).

**Table 3.** Average genotypic levels of kin response, nonkin response, kin recognition and nonkin invocation as evaluated by total root length (RL-TOTAL), given as percentages.

| **Cultivar name** | **Kin response** | **Nonkin response** | **Kin recognition** | **Nonkin invocation** |
| --- | --- | --- | --- | --- |
| Børsum * | -0.6% | 6.3% | 2.3% | 11.4% |
| Gammel Svensk Landhvede * | -5.7% | -0.5% | -0.8% | 6.2% |
| Lantvete från Dalarna * | 12.7% | 0.2% | 21.9% | 9.7% |
| Nordmøre * | 14.7% | 20.3% | -1.3% | 10.5% |
| Øland 5 * | 25.5% | 28.3% | 8.5% | 7.7% |
| Extra Squarehead | 4.7% | 6.3% | 0.6% | -2.2% |
| Vårpärl Svalöf | 6.1% | 8.1% | 2.4% | -1.3% |
| Tystofte Smaahvede | -1.5% | -4.3% | 8.1% | -7.9% |
| Als | -3.0% | 7.9% | -7.3% | -0.3% |
| Peragis | 6.4% | 11.8% | -1.4% | 2.4% |
| Extra Kolben II | 16.7% | 17.0% | 3.0% | -1.6% |
| Diamant | 1.5% | -0.8% | 8.0% | 3.6% |
| Atle | -3.7% | 15.2% | -15.0% | -10.2% |
| Progress | 10.9% | 4.8% | 10.2% | 8.9% |
| Zimmermanns** | -12.2% | -0.8% | -8.8% | -12.1% |
| Blanka | -15.4% | -12.8% | -2.3% | -9.5% |
| Touko | 9.1% | 8.5% | 4.0% | -5.0% |
| Rival | 4.3% | 15.3% | -6.7% | -2.8% |
| Vårpärl | 6.1% | -3.4% | 11.7% | -0.5% |
| Janus | 5.3% | 11.8% | -1.2% | 11.5% |
| Sappo | 1.9% | 4.1% | 3.0% | -1.4% |
| Saffran | 16.1% | 9.7% | 10.9% | 7.8% |
| Hja 21152 | -6.6% | 19.1% | -18.1% | 10.0% |
| William | 26.7% | 12.2% | 25.2% | 11.6% |
| Luja | 28.3% | 13.7% | 17.6% | 11.1% |
| Canon | 5.9% | 0.6% | 9.8% | 7.6% |
| Dragon | 8.9% | 13.1% | 2.0% | 11.5% |
| Curry | 2.4% | 9.7% | -3.5% | -5.0% |
| Fasan | 6.5% | 9.5% | 1.1% | -10.2% |
| *Average* | 5.9% | 8.0% | 2.9% | 2.1% |

\* Landrace
\*\* cv ‘Zimmermanns Begrannter Opferbaumer’

The landrace *Lantvete från Halland* showed clear signs of autotoxicity. For example, RL-TOTAL of this genotype was reduced by 44% in the KIN treatment compared to the PURE (control) treatment. In the gene bank registry, this accession is described as containing ‘different types with and without awn, white spike, coloured spike’. To avoid being unable to separate the effects of kin recognition and toxic allelopathy, this genotype was excluded from all analyses.

### Kin recognition and nonkin invocation

Comparison of root trait measurements in KIN treatments relative to NONKIN treatments presented a pattern in which kin recognition resulted in shorter, but not fewer roots (Table 2). Kin recognition differed among genotypes, particularly when evaluated based on RL-CV and RL-MAX; i.e. the interaction term *relatedness x genotype* was significant or marginally so (P = 0.019 and P = 0.087, respectively). The average kin recognition of receiver genotypes (across all tested nonkin donors) varied from positive to negative (as exemplified in Table 3) and did not change systematically with year of release (not shown).

When analysed combined as main factors in a linear model, both *donor* and *receiver* genotype were found to influence the root traits of the focal genotype (all P < 0.001, except the effect of *donor* on RL-CV with P < 0.01, and the effect of *donor* on RN, which was not significant). The same was found when accounting for relatedness as cofactor (not shown). When analysed in a diallel analysis of variance, the mean squares for effects of *donor* and *receiver (general kin effects, sensu* GCA; see statistics section) were found to be larger than those for specific combinations (*specific kin effects, sensu* SCA), especially for length and volume traits (Table 4). Donor and receiver genotypes generally explained a significant proportion of the observed variation in root traits, and highly significant mean squares for *reciprocals* showed that genotypes had different effect as *donor* than as *receiver* (Table 4).

**Table 4.**
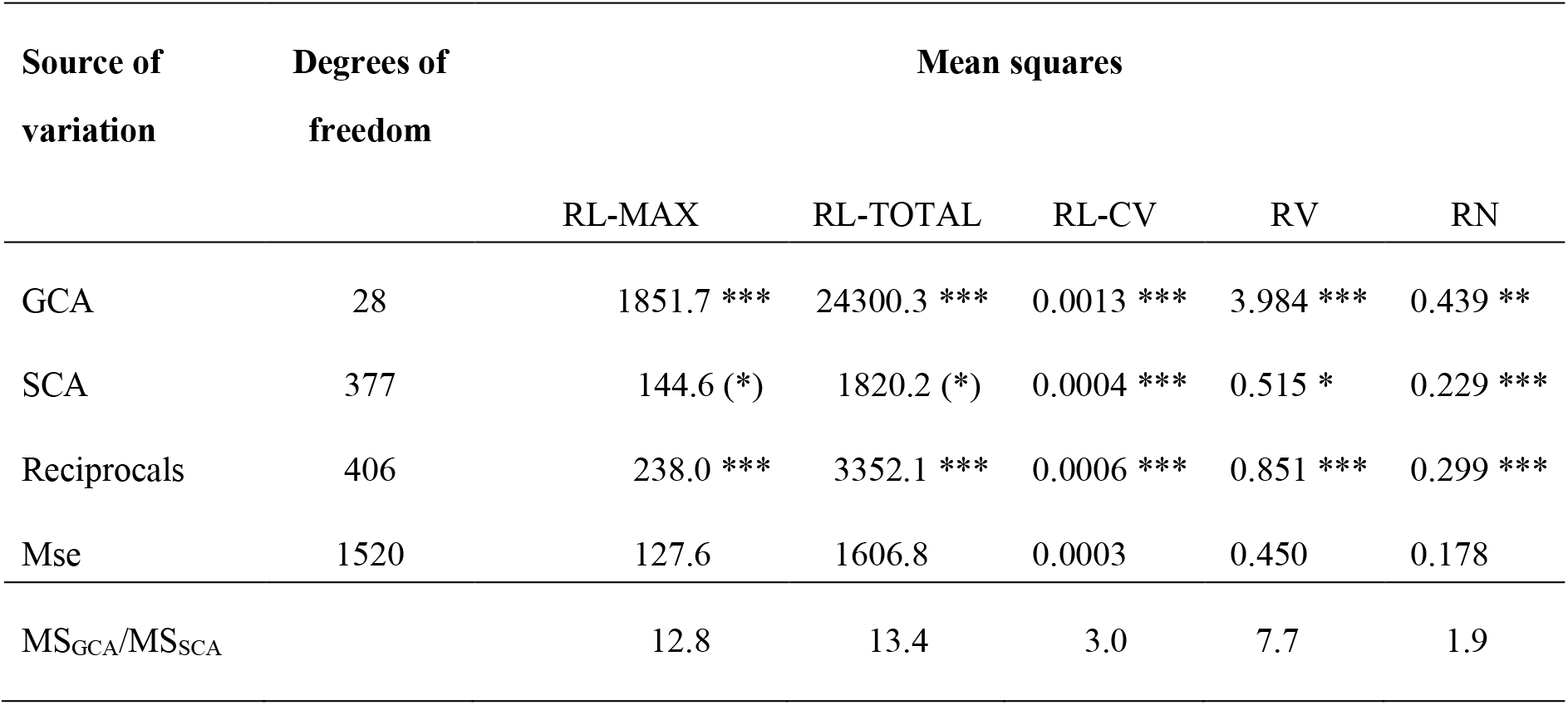
Summary of analysis of variance of the diallel setup with 29 genotypes acting as donors and receivers, analysed separately for each of five root traits. (*), *, **, *** denote marginal significance at 0.10 > *P* ≥ 0.05 and significance at *P* < 0.05, *P* < 0.01 and *P* < 0.001, respectively.

Average nonkin invocation of genotypes, i.e. their ability to invoke root trait response in other receiver genotypes relative to the kin responses of those receivers, varied from positive to negative for most root traits (as shown for RL-TOTAL in Table 3). However, RN showed predominantly negative levels of nonkin invocation, reflecting the generally positive kin responses for this root trait. The landrace showing signs of autotoxicity (*Lanthvete från Halland*) also produced exceptionally large nonkin invocation in the other genotypes for all traits (not shown), confirming the allelopathic effects of this genotype.

There were significant positive correlations between overall kin recognition and overall nonkin invocation of genotypes for each of the three root-length-related traits: genotypes showing higher levels of kin recognition also tended to invoke larger responses in other genotypes (Fig. 2). The five included landraces showed similar levels of kin recognition and nonkin invocation for all six root traits, predominantly invoking increased root length and volume across the set of receiver genotypes (Fig. 2a-c). The old cultivar *Vårpärl Svalöf* gave unusually positive nonkin invocation in root length variation (RL-CV) as compared to its kin recognition for this root trait. The Finnish cultivar *Hja 21152* had unusually negative RL-TOTAL in KIN treatment compared to its average across NONKIN treatments (seen as the upper left point in Fig. 2c). While this genotype had intermediate root length in the KIN treatment, it was the genotype with the longest roots across all NONKIN treatments.

**Figure 2.**
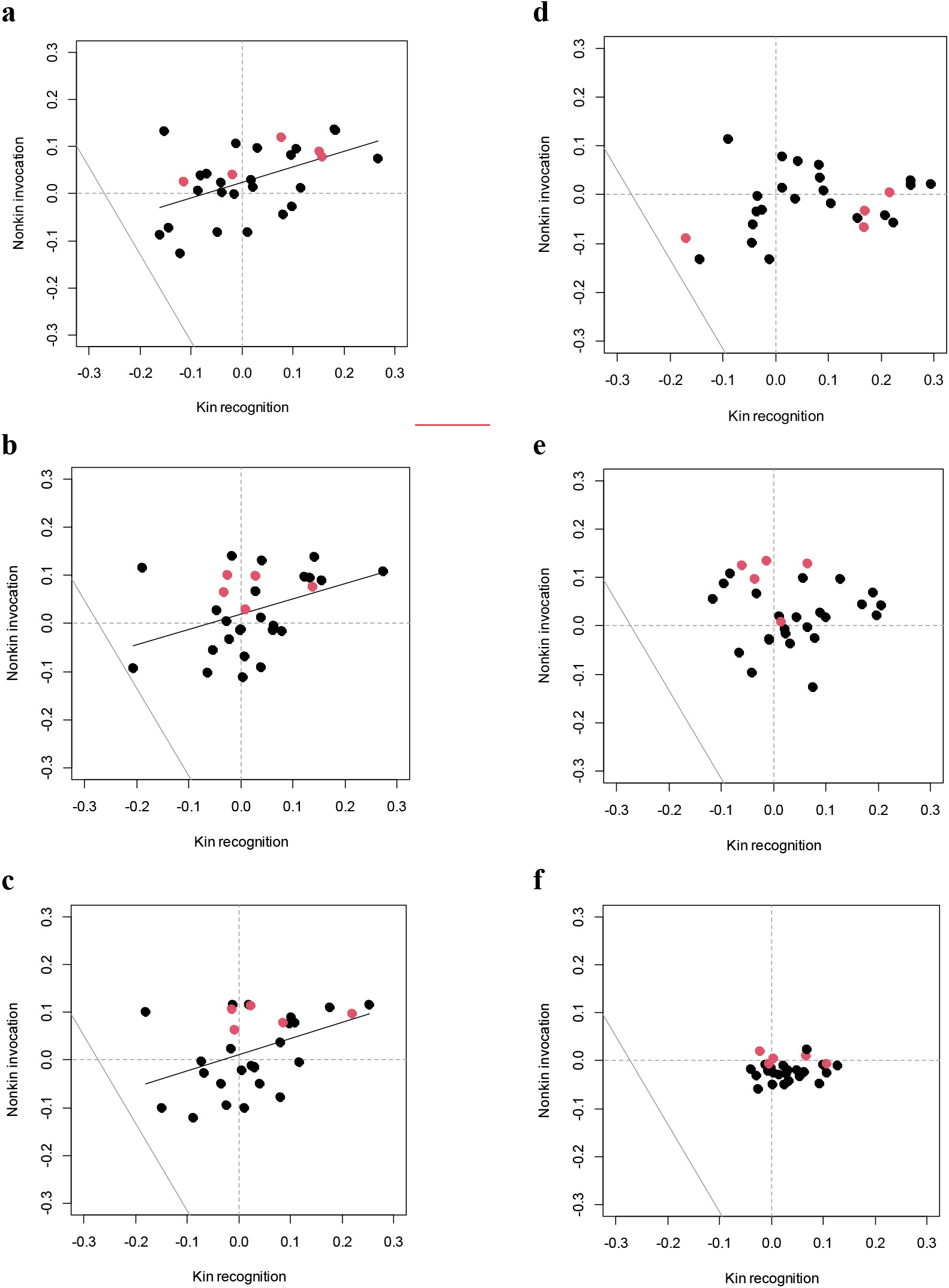
Relationships between overall kin recognition and nonkin invocation in (a) RL-MAX, (b) RL-PFS, (c) RL-TOTAL, (d) RL-CV, (e) RV and (f) RN. Red points denote the five included landraces. Full lines show significant linear regressions across all genotypes.

### Inhibitor effect

The 60 most responsive donor-receiver combinations were selected from the 30 combinations with the most positive levels of kin recognition and the 30 combinations with the most negative levels of kin recognition. These combinations were relatively evenly distributed across the involved genotypes, representing a total of 27 donor genotypes and 16 receiver genotypes. The two groups were analysed separately, each showing substantial and significant overall reductions in kin recognition in the presence of the chemical inhibitor, except for RL-CV among the combinations showing positive effects of kin recognition (Table 5).

**Table 5.** Tests for overall effect of chemical inhibitor on kin recognition in the most responsive genotype combinations (grouped into positive and negative kin recognition). ns, (*), *, ** and *** denote non-significance, marginal significance at 0.10 > P ≥ 0.05 and significance at P < 0.05, P < 0.01 and P < 0.001, respectively.

|  | Positive kin recognition |  |  |  |  | Negative kin recognition |  |  |  |  |
| --- | --- | --- | --- | --- | --- | --- | --- | --- | --- | --- |
|  | Control |  | Inhibitor |  | F-value | Control |  | Inhibitor |  | F-value |
| RL-MAX | 25% | *** | 4% | ns | 31.774 *** | -25% | *** | -4% | ns | 26.385 *** |
| RL-PFS | 33% | *** | 7% | (*) | 19.247 *** | -23% | *** | 0% | ns | 24.980 *** |
| RL-TOTAL | 32% | *** | 6% | ns | 20.749 *** | -21% | *** | 0% | ns | 17.640 *** |
| RL-CV | 25% | *** | 18% | ** | 0.721 ns | -28% | *** | -8% | * | 17.590 *** |
| RV | 29% | *** | 3% | ns | 24.267 *** | -18% | *** | 3% | ns | 14.015 *** |
| RN | 15% | *** | 3% | ns | 11.516 ** | -7% | *** | -2% | ns | 3.619 (*) |

## Discussion

The presented results support the hypothesis that wheat plants can distinguish kin from nonkin already in the earliest stages of growth and respond by changing their root growth pattern. Root length response to kin donors was generally lower than response to nonkin donors, aligning with kin selection theory and many previous studies (e.g. Semchenko et al. 2014).

Root growth was stimulated by preceding donors, whether these were kin or nonkin. The fact that these responses were higher in the longer roots suggests that root elongation was stimulated throughout the exposure period, with the longer roots being exposed for longer time. This corresponds to a model of root signalling in which the root tip, being the first plant part to explore new substrate, plays a crucial role in root responses to environmental stimuli (Doan et al. 2017, Sasse et al. 2018). Response to root neighbours, independent of relatedness, has been observed in rice (Fang et al. 2013). In that study, presence of kin neighbours resulted in reduced, not increased root length. In nature, outcrossing species such as rice are likely to face differently structured genetic neighborhoods than selfing species such as wheat, and it remains unanswered whether kin recognition behaviour generally differs between these reproductive groups of plants.

Kin recognition and nonkin invocation effects varied from negative to positive. Genotypes showing more positive kin recognition, responding more to kin than to nonkin substrate, generally also invoked stronger root growth in other genotypes. Similarly, genotypes growing shorter roots when exposed to kin compared to nonkin substrate also invoked shorter roots in other genotypes. This finding suggests that kin interaction is more complex than previously reported, while accommodating the reports showing variable responses or without evidence of kin recognition.

### Modes of indirect plant-plant interaction

Direct plant-plant interaction was made impossible by the experimental design. Genotypes could only affect each other indirectly via changes in the substrate environment. We identify four potential types of substrate change related to (i) the physical matrix, (ii) nutrient concentrations, (iii) presence of toxic compounds, and (iv) root exudates conferring kin recognition.

The donor seedling growing in a well could have caused physical changes to the substrate that may have affected the growth of the subsequent receiver seedling. The volume of substrate available to receivers was often observed to be visibly smaller than the volume of the originally dispensed substrate. This may have been due to some substrate sticking to the removed roots of the first seedling, despite efforts to leave all substrate in the well. Perhaps more likely, water uptake during donor seedling growth may have compressed the substrate matrix, reducing both the absolute and the relative water content. The expected effect of such a change would be reduced root growth in the receivers in KIN and NONKIN treatments, and hence, the general stimulation of receiver root growth suggests that this was not a dominant factor.

We assume that nutrient competition was not an important factor, given the short growth period in which seedlings are able to rely on seed nutrients, and the fact that the water agar solution contained practically no nutrients. Therefore, effects of niche partitioning (*sensu* File et al. 2012) are highly unlikely.

Exudation of toxic compounds by donor seedlings would be expected to impede *receiver* root growth. The landrace *Lanthvete från Halland* was excluded from the analyses as it clearly reduced the growth of receivers, indicative of toxicity. Some genotypes of wheat are known to produce allelochemicals supressing the growth in competing species (Wu et al. 2000), particularly benzoxazinoid hydroxamic acids (Niemeyer 2009). The findings of both positive and negative effects of kin and nonkin substrates on RL-TOTAL, compared to growth in pristine substrate, indicates that both toxic and kin recognition effects may have been in play. On the other hand, the average positive responses to kin and donors suggests that any toxic chemicals did not have major inhibitory effect on root growth.

Kin recognition was reduced by addition of the sodiumorthovanadate inhibitor. This supports the hypothesis that responses were largely due to *donor* release of chemical exudates to the substrate. We would not expect the inhibitor to moderate either the nutrient content or physical properties of the substrate, nor the response or seedlings to these environmental factors.

Recent studies have assessed kin response based on pot experiments, allowing simultaneous interaction (e.g. Fréville et al. 2019). This can be problematic as it is not possible to distinguish the effects of the indirect kin recognition from effects of more direct interaction such as competition for limited resources. Experiments that allow to study kin recognition effects until maturity without any direct interaction are difficult to design and involve other potentially confounding factors and trade-offs.

### Effects of relatedness

It is to date not clear how the degree of relatedness affects kin recognition in plants. One source of confusion has been that studies have used different definitions of kin and nonkin (the latter often called *stranger*), the former ranging from clonal ramets, over siblings, to members of the same population, and the latter ranging from non-sibling members of a population to individuals sampled from a distant population.

Here, seeds from the same cultivar was considered as kin, whereas seeds from other cultivars were considered as nonkin. This definition may be too broad for some cultivars if plants can only recognize full or half siblings. It is possible that the 10 earliest genotypes were not genetically pure, particularly the six landraces, yet, in any case it must be expected that what we call kin are more closely related to one another than to non-kin.

Based on our findings, we suggest that researchers of kin recognition need to study a wider range of genotypes with controlled levels of relatedness to establish (1) if kin recognition is a general phenomenon in plants, (2) the variability of kin responses within a set of genotypes, (3) what levels of relatedness plants are able to differentiate, and (4) the occurrence of specific vs. general kin recognition.

### Applied perspectives

The presented results clearly indicate that wheat can distinguish between kin and nonkin neighbours and that kin recognition exists also in modern varieties of bread wheat. It remains to be explored if and how kin recognition can contribute to the agronomic goal of maximizing total grain yield while reducing fertilizer requirements. In nature, kin recognition could help plants navigate complex environments, increasing fitness and promoting the survival of populations. Annual cropping systems, on the other hand, are characterised by a certain level of environmental control and distinctive fitness objectives somewhat different to those acting under natural selection.

The lack of systematic changes in kin recognition behaviour over the breeding period suggests that there has been no consistent selection on this trait, and that it is not correlated with other traits under selection. Meanwhile, it remains unknown if kin recognition could potentially interfere with water and nutrient acquisition (Finch et al. 2017). This would likely depend on the spatial response of root growth, i.e. any change in root architecture during kin response. There is recent evidence that breeding wheat for higher yields has generally resulted in fewer and deeper roots with less branching (Zhu et al. 2019), promoting uptake of water and nutrients from deeper soil layers as well as reduced inter-individual interaction. If targeted, breeding for root elongation mediated by kin recognition could support this trend even further.

In our experiments, seedlings were allowed to grow for a very short period. The observed increase in root length over the domestication period confirms the success of the common breeding strategy towards early establishment and growth, being decisive for the later plant biomass and competitive advantage over agricultural weeds. On the other hand, the observed kin recognition behaviour may not be representative for the effects over the whole lifespan of wheat plants. Furthermore, in vitro experiments such as ours leave out important elements likely to moderate chemical plant-soil and plant-plant interactions, including soil microorganisms and pedo-chemical processes.

### Conclusions

Based on the presented results, we propose that kin recognition be considered as a potential target for crop improvement to further promote crop soil foraging and reduce competitive interaction. This is particularly relevant for effective nutrient utilization under unfavourable conditions. Kin recognition ability in our field crops has potential to influence resource use efficiency of whole cropping systems, through altruistic sharing of soil resources, improved soil foraging and optimisation of investment in roots. Significant variation in kin recognition was found among earlier as well as later genotypes, ranging from positive to negative. This suggests that kin recognition is a quantitative trait determined by multiple genes, and that substantial genetic variation is available for this behaviour in wheat, also in more modern germplasm.

## Acknowledgements

The study was funded by The Danish Council for Independent Research, Technology and Production Sciences (FTP; grant no. 11-117112). The authors wish to thank Stina Christensen and Mads Nielsen for their help with root sampling and image analysis.

## References

Bais H.P. Shedding light on kin recognition response in plants. New Phytol 205, 4–6 (2015).

Bhatt M.V., Khandelwal A. & Dudley S.A. Kin recognition, not competitive interactions, predicts root allocation in young *Cakile edentula* seedling pairs. New Phytol 189, 1135–1142 (2011).

Biedrzycki M.L., Jilany T.A., Dudley S.A. & Bais H.P. Root exudates mediate kin recognition plants. Commun Integr Biol 3, 28–35 (2010).

Biedrzycki M.L., L V., Bais H.P. The role of ABC transporters in kin recognition in *Arabidopsis thaliana*. Plant Signal Behav 6, 1154–1161 (2011).

Biernaskie J.M. Evidence for competition and cooperation among climbing plants. Proc Biol Sci 278, 1989–1996 (2011).

Crepy M.A. & Casal J.J. Photoreceptor-mediated kin recognition in plants. New Phytol 205, 329–338 (2015).

Doan T.H., Doan T.A., Kangas M.J. et al. (2017). A low-cost imaging method for the temporal and spatial colorimetric detection of free amines on maize root surfaces. Front Plant Sci 8, 1513 (2017).

Donohue K. The Influence of Neighbor Relatedness on Multilevel Selection in the Great Lakes Sea Rocket. Am Nat 162, 1, 77–92 (2003).

Dudley S.A. & File A.L. Kin recognition in an annual plant. Biol Lett 3, 435–438 (2007).

Fang S., Clark R.T., Zheng Y. et al. Genotypic recognition and spatial responses by rice roots. Proc Natl Acad Sci USA 110, 2670–2675 (2013).

Finch J.A., Guillaume G., French S.A. et al. Wheat root length and not branching is altered in the presence of neighbours, including blackgrass. PLoS One 12, e0178176 (2017).

Fréville H., Roumet P., Rode N.O. et al. Preferential helping to relatives: A potential mechanism responsible for lower yield of crop variety mixtures? Evol Appl 12, 1837–1849 (2019).

Garnier E., Navas M.-L., Grigulis K. Trait-based ecology: definitions, methods, and a conceptual framework (Chapter 2), in: Plant Functional Diversity: Organism traits, community structure, and ecosystem properties. pp. 9–25 (2015).

Hamilton W.D. The genetical evolution of social behaviour. I J Theor Biol 7, 1–16 (1964).

Lepik A., Abakumova M., Zobel K. & Semchenko M. Kin recognition is density-dependent and uncommon among temperate grassland plants. Funct Ecol 26, 1214–1220 (2012).

Li J., Xu X. & Feng R. Soil fertility and heavy metal pollution (Pb and Cd) alter kin interaction of *Sorghum vulgare*. Environ Exp Bot 155, 368–377 (2018).

Marler T.E. Kin recognition alters root and whole plant growth of split-root *Cycas edentata* seedlings. Hort Sci 48 (10), 1266–1269 (2013).

Masclaux F. et al. Competitive ability not kinship affects growth of *Arabidopsis thaliana* accessions. New Phytol 185, 322–331 (2010).

McMickle G.G. Brown J.S. An ideal free distribution explains the root production of plants that do not engage in a tragedy of the commons game. J Ecol 102, 963–971 (2014).

Milla R., Forero D.M., Escudero A., Iriondo J.M. Growing with siblings: a common ground for cooperation or for fiercer competition among plants? Proc R Soc B 276, 2531–2540 (2009).

Monzeglio U. & Stoll P. Effects of spatial pattern and relatedness in an experimental plant community. Evol Ecol 22, 723–741 (2008).

Murphy G.P. & Dudley S.A. Kin recognition: Competition and cooperation in *Impatiens* (Balsaminaceae). Am J Bot 96, 1990–1996 (2009).

Niemeyer H.M. Hydroxamic acids derived from 2-hydroxy-2H-1,4-benzoxazin-3(4H)-one: key defense chemicals of cereals. J Agric Food Chem 57, 1677–1696 (2009).

R Core Team (2020). R: A language and environment for statistical computing. R Foundation for Statistical Computing, Vienna, Austria. URL: https://www.R-project.org/.

Salamini F., Özkan H., Brandolini A. et al. Genetics and geography of wild cereal domestication in the near east. Nat Rev Genet 3, 429–441 (2002).

Sasse J., Martinoia E., Northen T. (2018). Feed your friends: do plant exudates shape the root microbiome? Trends Plant Sci. 23, 25–41.

Sattler J. & Bartelheimer M. Root responses to legume plants integrate information on nitrogen availability and neighbour identity. Basic Appl Ecol 27, 51–60 (2018).

Semchenko M., Saar S. & Lepik A. Plant root exudates mediate neighbour recognition and trigger complex behavioural changes. New Phytol 204, 631–637 (2014).

Weiner J. Allocation, plasticity and allometry in plants. Perspect. Plant Ecol Evol Syst 6, 207–215 (2004).

Wu H., Pratley J., Lemerle D. & Haig T. Evaluation of seedling allelopathy in 453 wheat *(Triticum aestivum*) accessions against annual ryegrass (*Lolium rigidum*) by the equal-compartment-agar method. Aust J Agric Res 51, 937 (2000).

Yaseen M. DiallelAnalysisR: Diallel Analysis with R. R package version 0.1.1. URL: https://CRAN.R-project.org/package=DiallelAnalysisR (2016).

Yee T.W. (2020). *VGAM:* Vector Generalized Linear and Additive Models. R package version 1.13. URL: https://CRAN.R-project.org/package=VGAM.

Zhu Y.H., Weiner J., Yu M.X., Li F.M. Evolutionary agroecology: Trends in root architecture during wheat breeding. Evol Appl 12 (4), 733–743 (2019).

