## Supplementary tables and figures for "Effects of kin recognition on root traits of wheat germplasm over 100 years of breeding"

### Supplementary material

**Table S1**. Information on gene bank accession number, country of origin, year of release, landrace status and seasonality of all genotypes included in the study.

| Genotype | Gene bank accession no. | Country of origin | Year of release [b] |
|---|---|---|---|
| Børsum | NGB2125 | Norway | LR |
| Gammel Svensk Landhvede * | | Sweden | LR |
| Lantvete från Dalarna | NGB8199 | Sweden | LR |
| Lantvete från Halland | NGB11080 | Sweden | LR |
| Nordmøre | NGB6673 | Norway | LR |
| Øland 5 * | NGB6674 | Sweden | LR |
| Extra Squarehead * | NGB6694 | Great Britain | 1900 |
| Vårpärl Svalöf | | Sweden | 1901 |
| Tystofte Smaahvede * | NGB13446 | Denmark | 1909 |
| Als * | NGB4770 | Denmark | 1923 |
| Peragis | TRI 559 | Germany | 1923 |
| Extra Kolben II | NGB8923 | Sweden | 1926 |
| Diamant | NGB6679 | Sweden | 1928 |
| Atle | NGB7455 | Sweden | 1936 |
| Progress | NGB6411 | Sweden | 1941 |
| Zimmermanns [a] | TRI 836 | Germany | 1949 |
| Blanka | NGB9691 | Sweden | 1950 |
| Touko | NGB359 | Finland | 1950 |
| Rival | NGB6684 | Sweden | 1952 |
| Vårpärl | NGB6675 | Sweden | 1959 |
| Janus | TRI 28644 | Germany | 1967 |
| Sappo | NGB7467 | Sweden | 1971 |
| Saffran | NGB7472 | Sweden | 1978 |
| Hja 21152 | NGB350 | Finland | 1979 |
| William | NGB7059 | Sweden | 1979 |
| Luja | NGB357 | Finland | 1981 |
| Canon | NGB7481 | Sweden | 1988 |
| Dragon | NGB9954 | Sweden | 1988 |
| Curry | NGB11708 | Sweden | 1994 |
| Fasan | NGB9675 | Germany | 1997 |

* Winter type; all others are spring type

[a] cv 'Zimmermanns Begrannter Opferbaumer'

[b] LR are landraces with no available release year (presumed 19$^{th}$ century)

**Table S2**. Basic root traits of the studied genotypes grown in pristine growth medium, including length of the longest root (RL-MAX), total root length (RL-TOTAL), total root volume (RV), total root length of the five longest roots (PFS) and total number of roots (RN).

| Genotype | RL-MAX (cm) | RL-PFS (cm) | RL-TOTAL (cm) | RL-CV (%) | RV ($mm^3$) | RN |
|---|---|---|---|---|---|---|
| Børsum | 5.94 | 21.65 | 22.5 | 5.8 | 24.6 | 5.6 |
| Gammel Svensk Landhvede | 5.99 | 18.98 | 19.2 | 9.4 | 17.8 | 5.2 |
| Lantvete från Dalarna | 6.34 | 22.16 | 22.4 | 8.1 | 34.7 | 5.2 |
| Lantvete från Halland | 6.44 | 25.03 | 25.9 | 6.2 | 49.0 | 5.4 |
| Nordmøre | 4.59 | 18.01 | 17.2 | 6.3 | 19.2 | 5.0 |
| Øland 5 | 6.84 | 20.81 | 22.1 | 8.4 | 27.7 | 5.4 |
| Extra Squarehead | 5.54 | 20.41 | 21.0 | 5.9 | 27.9 | 5.4 |
| Vårpärl Svalöf | 8.01 | 27.40 | 25.7 | 8.2 | 22.7 | 4.6 |
| Tystofte Smaahvede | 6.39 | 22.60 | 20.9 | 9.1 | 29.4 | 4.6 |
| Als | 6.13 | 19.40 | 19.3 | 7.8 | 25.1 | 4.8 |
| Peragis | 6.21 | 22.95 | 23.1 | 6.6 | 34.3 | 5.3 |
| Extra Kolben II | 6.64 | 22.75 | 23.5 | 6.6 | 24.6 | 5.3 |
| Diamant | 5.27 | 18.28 | 18.7 | 7.9 | 25.4 | 5.4 |
| Atle | 6.07 | 22.43 | 22.9 | 6.9 | 36.1 | 5.5 |
| Progress | 6.88 | 23.33 | 24.0 | 7.6 | 28.1 | 5.5 |
| Zimmermanns | 6.81 | 23.82 | 24.3 | 8.0 | 37.0 | 5.5 |
| Blanka | 6.69 | 24.84 | 24.7 | 6.5 | 29.7 | 5.0 |
| Touko | 5.82 | 21.09 | 22.1 | 6.4 | 32.6 | 5.6 |
| Rival | 6.72 | 23.14 | 22.6 | 7.6 | 33.4 | 4.9 |
| Vårpärl | 7.78 | 27.04 | 26.9 | 7.7 | 33.2 | 5.1 |
| Janus | 5.89 | 21.25 | 21.1 | 6.0 | 42.7 | 5.0 |
| Sappo | 6.94 | 25.49 | 26.6 | 7.1 | 43.5 | 5.6 |
| Saffran | 4.67 | 16.84 | 16.8 | 7.2 | 26.4 | 5.1 |
| Hja 21152 | 7.71 | 26.37 | 27.3 | 7.2 | 39.2 | 5.4 |
| William | 5.75 | 21.64 | 21.4 | 7.6 | 33.7 | 5.3 |
| Luja | 8.13 | 25.53 | 26.5 | 9.3 | 35.6 | 5.5 |
| Canon | 8.31 | 27.00 | 26.9 | 9.1 | 39.0 | 5.2 |
| Dragon | 7.20 | 25.87 | 26.7 | 6.8 | 45.5 | 5.4 |
| Curry | 6.82 | 22.93 | 23.7 | 9.2 | 34.5 | 5.5 |
| Fasan | 6.09 | 19.70 | 20.4 | 8.2 | 33.8 | 5.1 |

**Figure S1.** Example of (a) an original root scan and (b) the resulting processed image (white pixels belonging to one of the identified roots).

**A**

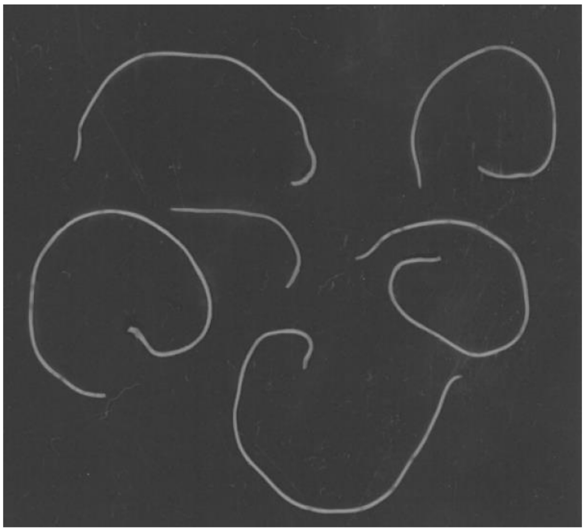

**b**

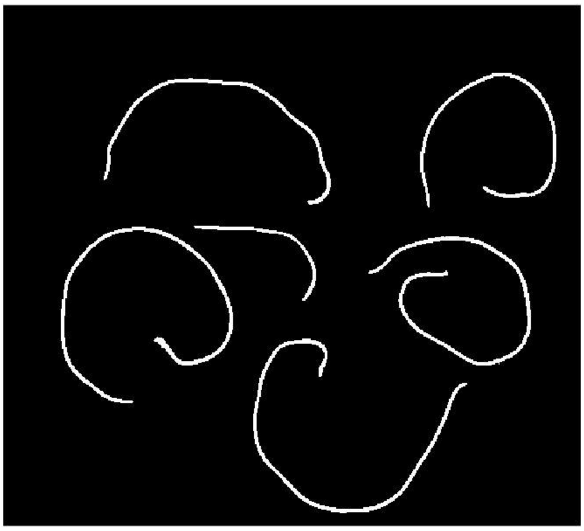

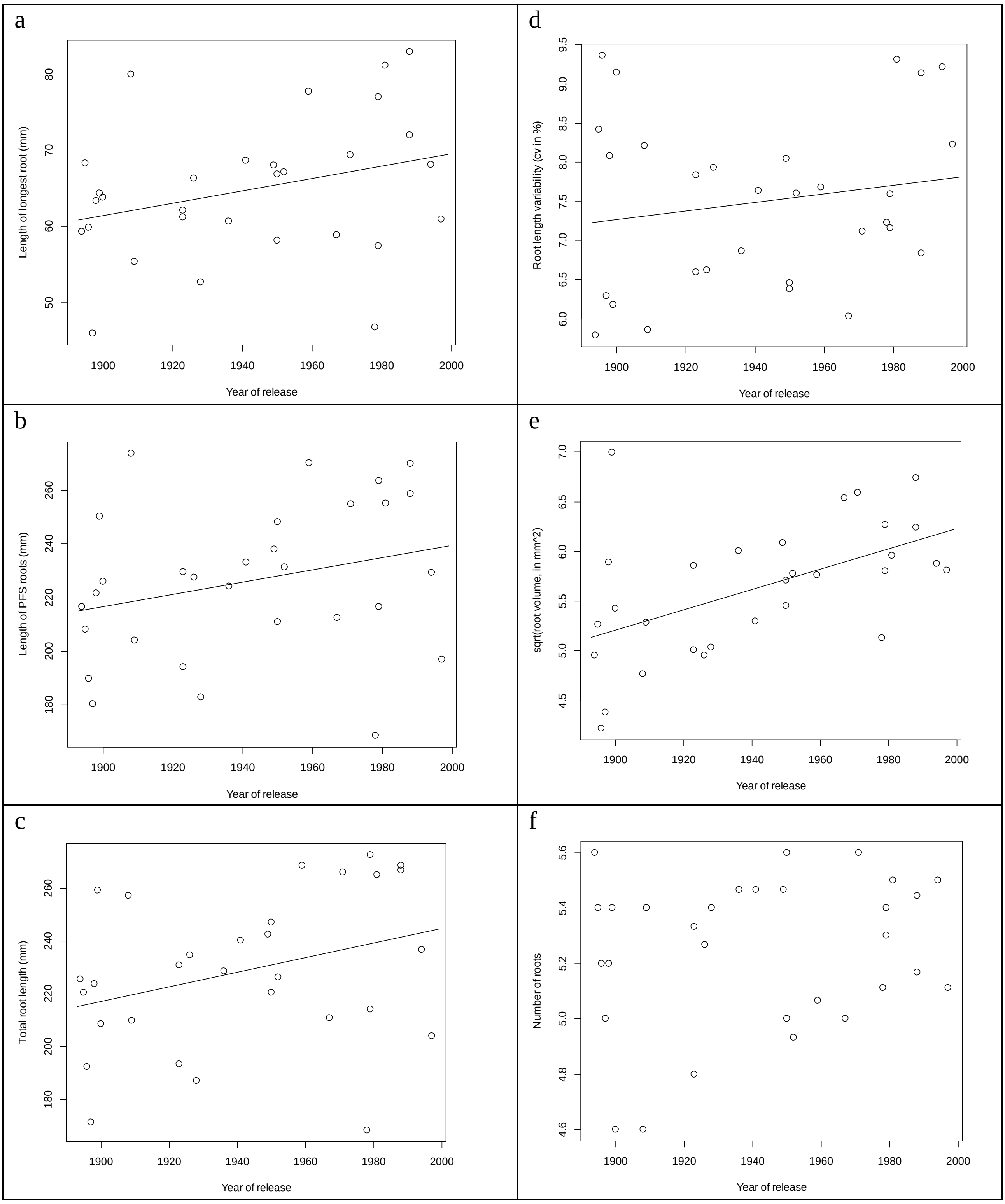


**Fig S2**. Root traits of genotypes when grown in pristine substrate (PURE), as changing over the studied release period (landraces were assigned a release year, as described in the main text). Full lines show the significant linear regressions of RL-MAX (a), RL-PFS (b), RL-TOTAL (c), RL-CV (d), RV (e) and RN (f).
